# Spatial gene-by-environment mapping for schizophrenia reveals locale of upbringing effects beyond urban-rural differences

**DOI:** 10.1101/315820

**Authors:** Chun Chieh Fan, John J. McGrath, Vivek Appadurai, Alfonso Buil, Michael J. Gandal, Andrew J. Schork, Preben Bo Mortensen, Esben Agerbo, Sandy A. Geschwind, Daniel Geschwind, Thomas Werge, Wesley K. Thompson, Carsten Bøcker Pedersen

## Abstract

Identification of mechanisms underlying the incidence of psychiatric disorders has been hampered by the difficulty in discovering highly-predictive environmental risk factors. For example, prior efforts have failed to establish environmental effects predicting geospatial clustering of schizophrenia incidence beyond urban-rural differences. Here, we employ a novel statistical framework for decomposing the geospatial risk for schizophrenia based on *locale of upbringing* (place of residence, ages 0-7 years) and its synergistic effects with *genetic liabilities* (polygenic risk for schizophrenia). We use this statistical framework to analyze unprecedented geolocation and genotyping data in a case-cohort study of n=24,028 subjects, drawn from the 1.47 million Danish persons born between 1981 and 2005. Using this framework we estimate the effects of upbringing locale (E) and gene-by-locale interactions (GxE). After controlling for potential confounding variables, upbringing at high-risk locales increases the risk for schizophrenia on average by 122%, while GxE modulates genetic risk for schizophrenia on average by 78%. Within the boundaries of Copenhagen (the largest and most densely populated city of Denmark) specific locales vary substantially in their E and GxE effects, with hazard ratios ranging from 0.26 to 9.26 for E and from 0.20 to 5.95 for GxE. This study provides insight into the degree of geospatial clustering of schizophrenia risk, and our novel analytic procedure provides a framework for decomposing variation in geospatial risk into G, E, and GxE components.

## Introduction

For public mental health, it is critical to know which environmental factors can be modified to mitigate the risk of psychiatric disorders. However, identifying modifiable environmental factors has been a contentious issue^1,2^, especially when the effects may depend on one’s genetic liability for illness. Take as an example one of the best-established environmental risks for schizophrenia, childhood upbringing in an urban area. Persons born and raised in urban areas have an approximately two-fold increased risk of schizophrenia compared to those born and raised in rural areas ^3,4^. Researchers have rigorously examined potentially causal elements of urban upbringing, such as accessibility to health care ^3,5^, selective migration of individuals ^6^, airpollution ^7^, infections ^8^, and socioeconomic inequality ^9^. None of these candidate elements explain a substantial amount of the urbanicity-related increase in risk ^3,6-9^. The conditional relationship between genetic liabilities and environmental factors are even harder to detect despite some cohort studies suggesting an interaction between urban upbringing and family history of schizophrenia ^10-13^.

The difficulty in isolating specific environmental risk elements underlying urbanicity effects on schizophrenia incidence exemplifies a serious methodological challenge. The process for discovering environmental risk factors typically relies on a hypothesis-driven “candidate environmental factor” approach. Researchers need to formulate a carefully-constructed environmental hypothesis, measure it, and then determine if it associates with risk of the disease, usually in a study of selected participants not necessarily representative of the entire population of interest. Similar to the candidate gene approach before the dawning of genome-wide association studies (GWAS) ^14^, the candidate environment approach suffers from the “spotlight effect”, ignoring the likely complexity of many environmental factors interacting with each other and with genetic liabilities to determine overall risk for illness. Even with a plausible hypothesis, measurement of the specific environmental factor may be imprecise, masking its relationship to the illness. For example, social epidemiologists rely on instruments as proxy measures for socioeconomic inequalities. Given the complexity of real-life socio-economic forces, lack of association with schizophrenia could be caused by instrument measurement error or because the instrument does not capture the relevant social-economic factors ^9^.

An alternative to the candidate environment approach is to assess spatial patterns of disease risk without directly measuring environmental factors. As with John Snow isolating the environmental source of cholera outbreak via mapping the cases ^15^, identifying spatially-localized disease “hot spots” can assist in the discovery of latent environmental factors. Advanced methods for disease mapping have been developed within the field of geostatistics, particularly in applying spatial random effect models to infer latent environmental variation in causal risk factors ^16^. As the urbanicity-related increase in risk for schizophrenia was first noted through spatial clustering of disease incidence ^17^, inferring risk hot spots to a finer resolution may provide insight into potential risk-modulating environmental elements before investing substantial resources in active measurement.

With this concept in mind, we develop a disease mapping strategy to address the need for discovering environmental factors without direct measurement. We use spatial random effects to map the geographic distribution of genetic liabilities (G), locale of upbringing (E), and their synergistic effects (GxE) on disease risk. By treating E and GxE as latent random fields, we avoid the methodological biases inherent in the candidate environment approach. As a proof of concept, we examine geospatial variation in schizophrenia risk across Denmark. To do so, we apply this novel analytical approach to data from an unprecedented, population-based case-cohort study that includes subject genotyping and detailed residential information from birth up to age seven years. We are thus able to assess locale of upbringing effects on schizophrenia risk with a resolution beyond conventionally-defined levels of urbanicity, allowing us to assess variation in spatial risk, and to ask whether spatially-localized environmental factors modulate genetic liability of risk for schizophrenia.

## Method

Our spatial mapping approach follows three steps: 1) defining neighboring locales to characterize the latent environment field, 2) estimating random effects associated with each locale, and 3) mapping the spatial distribution based on the realized effects on locales. These three steps are calibrated to ensure a good balance between fine spatial resolution and adequate statistical power. Furthermore, the modeling strategy partitions observed effects on risk for schizophrenia into different components: locale of upbringing (E), genetics (G), and the synergistic effects of spatial locale and genetics (GxE).

## Definition of locales

We exploit the duality between Delaunay triangulation and Voronoi tessellation^18^, ensuring each defined locale has a sufficient number of study subjects without biasing resulting locales towards metropolitan regions. This balance is important, as some metropolitan areas have more than 100 study subjects living in the same 1 km^2^ region, while some rural areas have only one subject, with no neighboring subjects within a several km radius. The algorithm achieves the balance between resolution and sufficient number of samples in each locale by adaptively merging neighboring locales with too few individuals into larger locales (Figure S1).

The primary advantage obtained from achieving a balance between resolution and number of individuals in each locale is to localize the regions as much as possible while retaining high statistical power to estimate locale (E) and gene x locale (GxE) spatial random effects. As a byproduct, the population density of each locale is also automatically calculated from this procedure, since the area of each locale is inversely proportional to the population density. Consistent with previous studies on urbanicity-related schizophrenia risk ^3^, we use the derived population density as a surrogate measure of urbanicity in subsequent analyses.

## Estimating the effects associate with the locale

Mixed effects models provide the necessary tools to estimate the latent environmental and gene x environmental effects. Fixed effects in the model control for potential confounding factors, whereas random locale effects approximate the latent field across all spatial locations. Once the random effect variance is estimated and determined to be significantly greater than zero, spatial mapping is achieved through computing the posterior means of the random effects for each locale, defined by the best linear unbiased predictors ^16^.

To ensure the validity of this approach, we performed 1000 Monte Carlo simulations to determine how well we can estimate E and GxE via the spatial mixed effects model. Given a sample size of 30,000 individuals with disease prevalence of one percent and heritability of 70 percent (similar to the profile of schizophrenia), we obtain an unbiased estimation of spatial effects (E), while GxE effects are conservatively bounded by the predictive power of the genetic instrument (Figure S2 and Supplemental Information).

## Empirical Study

We demonstrate the feasibility of our spatial mapping approach by characterizing E and GxE effects of schizophrenia in the Danish population. To map the synergistic effects of locale of upbringing and schizophrenia genetic liability, chronological residential information and genotyping data from the same population-based cohort is needed. The Danish Lundbeck Foundation Initiative for Integrative Psychiatric Research (iPSYCH) case-cohort study provides a unique opportunity for this aim^19^. Prior to iPSYCH, genome-wide association studies (GWAS) of psychiatric disorders have lacked information on locale of upbringing, while population registry studies with detailed residential locales have not yet implemented polygenic data analyses. By linking with the Danish Civil Registration System, iPSYCH has a nationally-representative sample with whole-genome genotyping and detailed chronological residential information. Together with the case-cohort design^17^, these characteristics of iPSYCH enable us to obtain nationally representative estimates of the locale effects and the modulating effects of locale on genetic risk.

For this analysis, we extracted genotyped schizophrenia cases and a population random sample cohort from the iPSYCH study ^19^. The study design and sample description has been published elsewhere ^19^. A flow chart of the recruitment can be found in the Supplementary Information (Figure S3). In short, cohort members (N = 30,000) were randomly sampled individuals from the entire Danish population born between 1981 and 2005 and surviving past one year of age (N = 1,472,262). Cases with schizophrenia were ascertained through the Danish Psychiatric Central Research Register, using diagnostic classifications based on the International Classification of Diseases, 10th revision, Diagnostic Criteria for Research (Diagnostic code F20; ICD-10-DCR). Patients with schizoaffective disorders were excluded. All psychiatric contacts until December 31, 2013 were obtained from the register, resulting in 3,540 genotyped individuals diagnosed with schizophrenia. The residential locations of case-cohort members were obtained through linkage to the Danish Civil Registration System. To focus on the early life experience, i.e. upbringing effects, the residential location of an individual was retrieved at three ages: at birth, age 5 years, and age 7 years. Individuals’ exact locations were blurred to 1 km^2^ grid cells to protect privacy. DNA samples were obtained from the Danish Neonatal Screening Biobank and sequenced with Infinium PsychChip v1.0 array (Illumina, San Diego, California, United States of America).

To prevent confounds due to recent migration and large-scale ethnic differences, we restrict our analyses to unrelated individuals who are of European descent, as determined by genetic ancestry^20,21^ and with both parents born in Denmark based on Danish registry information. The final analyses include 24,028 case-cohort members (2,328 schizophrenia cases, 21,700 cohort members) who met above criteria and passed genotyping quality controls. Table S1 demonstrates the basic demographic characteristics of the included case-cohort.

## Genotype Processing and Deriving Polygenic Risk Scores

Eleven million single nucleotide polymorphisms (SNPs) were imputed based on genotyped SNPs that pass the following criteria: minor allele frequencies greater than 1 percent; frequencies in Hardy-Weinberg Equilibrium; SNPs autosomal and bi-allelic. Genetic principal components were derived based on quality controlled genotyped SNPs using flashPCA^21^. SHAPEIT3 was used for phasing^22^ and IMPUTE2 was used for imputation^23^. The reference panel was 1000 genomes project phase 3 ^24^.

To obtain a genetic instrument with good predictive power for detecting GxE, we calculated the polygenic risk score (PRS) using the summary statistics for 34,129 cases and 45,512 controls from the Psychiatric Genomics Consortium (PGC) Schizophrenia GWAS^25^. The PRS is the sum of the products of effect sizes of SNPs estimated from this independent GWAS and the dosage of those SNPs from the iPSYCH case-cohort. The included SNPs were pruned to ensure independence, while no significance threshold was set to filter SNPs. Parameters for calculating PRS include clumping (r^2^ = 0·1, distance = 250kb), and pruning (r2 = 0·8, window = 2kb, increment = 2kb). Nonetheless, PRS is inherently a weak genetic instrument, so our estimate on GxE should be regard as a conservative lower bound of interaction effects (Figure S2).

## Code Availability

The code used for simulations, empirical analysis, and visualization can be found at (https://chunchiehfan.shinyapps.io/iPSYCH_geo_tess_SZ/). The interactive version of the disease mapping is shown on the web portal while all the relevant codes can be downloaded on it. All analyses are implemented in R^26^. R packages employed include spatstat^27^ and coxme^28^. The geographical visualization is done with ggmap ^29^, which extracts geographical information from Google Maps. An interactive version of the risk map is generated using leaflet ^30^ and shiny ^31^.

## Results

### Spatial distribution of overall risk of schizophrenia

We utilize the entire population cohort of iPSYCH, excluding cases, to derive locales. The resulting map contains 186 non-overlapping locales, with the number of cohort members ranging from 65 to 197 individuals in each locale (median = 105). To demonstrate the risk distribution across defined locales, we use the Mantel-Haenszel approach for estimating risk ratios (RR) while correcting for age differences ^32^. Figure 1 displays the RR’s from the Mantel-Haenszel analyses. With the exception of the southwestern portion of Denmark, the majority of rural regions have lower risk ratios while high-risk locales are concentrated in large cities (Figure 1a). By plotting RR’s against the size of each locale, Figure 1b demonstrates a general trend for spatial risks of schizophrenia, meaning locales with higher population density tend to have higher RR’s. Thus, the risk distribution recapitulates the known urbanicity effects. However, there is substantial variation in risk even controlling for locale size; for example, RR’s can range from protective to highly detrimental within densely populated areas (Figure 1b).

**Figure 1.**
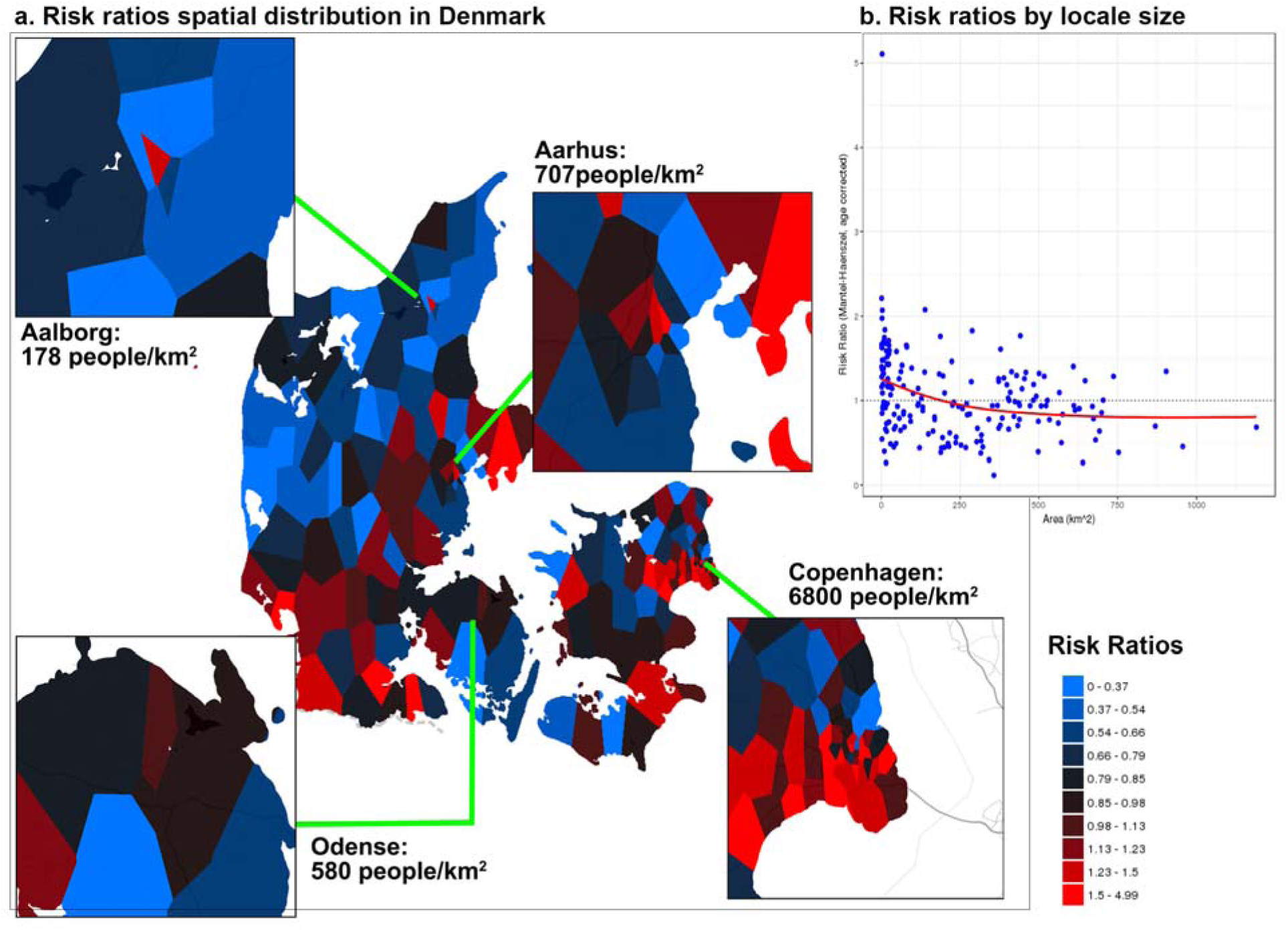
Age corrected risk ratios of schizophrenia for each locale comparing to national average. The risks are estimated based on case-cohort counts stratified by age, as Mantel-Haenszel estimates. (a.) Mantel-Haenszel estimated RR for Denmark. Four largest cities were further zoomed in as their corresponding population densities were annotated below. The metropolitan area (Copenhagen) has highest population density and also clusters of high-risk areas. Lower the population density tends to have lower disease risk except the regions such as western-southern region of Denmark. For visualization purpose, colors were centered on one as national average while scaled according to risk deciles. (b.) RR of each locale is plot against the associated size of locale. The dots represent each locale while the red solid line is the overall trend based on smoothed spline.

### The Contribution of the E and GxE

Next, we implement the spatial mixed effects model to identify sources of variation in the observed risk across locales. Given the concern of potential confounds, all models include fixed effects of gender, the first three genetic principal components, and family history as covariates. Survival models were used to account for age distribution ^19^ and observations were weighted by the inverse of each subject’s sampling probability ^33^ for inclusion in iPSYCH. Time-to-event is defined as age at first hospital contact for schizophrenia for cases, and the minimum of age of death, disappearance, immigration, or age at date of registry information collection (December 31, 2013) for cohort members without schizophrenia. For comparison purposes, we also fit a model with fixed effect of the covariates and no random effects (Model 1). Results are shown in Table 1.

**Table 1.** Hazard ratios estimates from four nested models based on iPSYCH case-cohort

|  | Model 1 |  | Model 2 |  | Model 3* |  |  |
| --- | --- | --- | --- | --- | --- | --- | --- |
|  | HR | 95%CI | HR | 95%CI | HR | 95%CI | p-value |
| Individual Level |  |  |  |  |  |  |  |
| Gender (Male) | 1.05 | (0.99 - 1.11) | 1.06 | (1.00 - 1.11) | 1.08 | (1.01 - 1.13) | 0.01 |
| Genetic PC 1 | 1.07 | (1.04 - 1.10) | 1.08 | (1.05 - 1.12) | 1.15 | (1.11 - 1.18) | 2x10<sup>-15</sup> |
| Genetic PC 2 | 0.92 | (0.89 - 0.94) | 0.97 | (0.95 - 1.01) | 0.99 | (0.96 - 1.02) | 0.59 |
| Genetic PC 3 | 0.92 | (0.89 - 0.95) | 0.92 | (0.90 - 0.95) | 0.93 | (0.90 - 0.95) | 4x10<sup>-7</sup> |
| Family History <sup>1</sup> | 6.07 | (5.23 - 7.05) | 4.61 | (3.93 - 5.04) | 5.63 | (4.75 - 6.67) | <2x10<sup>-16</sup> |
| PRS <sup>2</sup> | 1.27 | (1.24 - 1.31) | 1.26 | (1.23 - 1.29) | 1.34 | (1.21 - 1.49) | 2x10<sup>-8</sup> |
| Spatial Level |  |  |  |  |  |  |  |
| Population Density (Urban vs Rural) <sup>3</sup> | 1.89 | (1.53 - 2.33) | 1.64 | (0.51 - 5.23) | 1.46 | (0.49 - 4.38) | 0.49 |
| E <sup>4</sup> |  |  | 2.29 |  | 2.22 |  | <2x10<sup>-16</sup> |
| GxE <sup>4</sup> |  |  |  |  | 1.78 |  | <2x10<sup>-16</sup> |

1. Either parent has been diagnosed with schizophrenia, based on the national psychiatric registry 2. PRS has been zero centered and standardized to unit variance 3. Unit increase corresponds to going from 55 person/km<sup>2</sup> to 5220 person/km<sup>2</sup>, equivalent to previous definition of rural to urban residence. 4. The hazard ratios for E and GxE are median hazard ratios (median of hazard ratio absolute difference over all possible pairs of regions). P values are based on likelihood ratio test to the model without random effects.
\* Model 3 is a full interaction model, obtained by multiplying PRS with all other covariates. Since E and GxE are the effects of interest, no other interactions are shown here. P-values shown are for Model 3.

Compared to rural regions, being born and living in densely populated urban area increases the risk of schizophrenia by (hazard ratio = 1.89, 95% CI: 1.53 - 2.33), which replicates previous studies on urbanicity effects ^3,4^. The inclusion of spatial random effects (E) reduces the urbanicity effect to hazard ratio = 1.64 with confidence interval encompassing 1. Model 3 with both E and GxE effects significantly contributes explanatory power to the variation in risk for schizophrenia (Log-likelihood ratio tests p < 2×10^−16^), while the urbanicity effect is further reduced (hazard ratio = 1.46). Model 3 already contains interaction terms of variables included in the model, hence it is unlikely that GxE effects can be attributed to confounding interactions ^1^. Median hazard ratios for E and GxE components, defined as the median absolute difference in hazard ratios for all possible combinations of pairs of locales ^34^, are 2.22 and 1.78, respectively, representing a 122% and 78% expected change in risk if living in a high-risk locale.

### Spatial distribution of the risk components of schizophrenia

The geographical distribution of E and GxE are shown in Figure 2. The E component mirrors the heightened risk in the southwestern part of the Denmark (Figure 2a) and the southern portion of Copenhagen, the metropolitan area with highest population density (Figure 2b). However, within the city boundary, hazard ratios vary strongly from protective to highly detrimental (hazard ratio: 0.26 to 9.26, Figure 2c). The GxE component has a different spatial pattern compared to E (Figure 2d). Within the metropolitan boundary, high-risk GxE locales are concentrated in the city center (Figure 2e) and the modulating effect can range from a decrease of risk of 80% to a 6-fold increase (hazard ratios: 0.20 to 5.95, Figure 2f).

**Figure 2.**
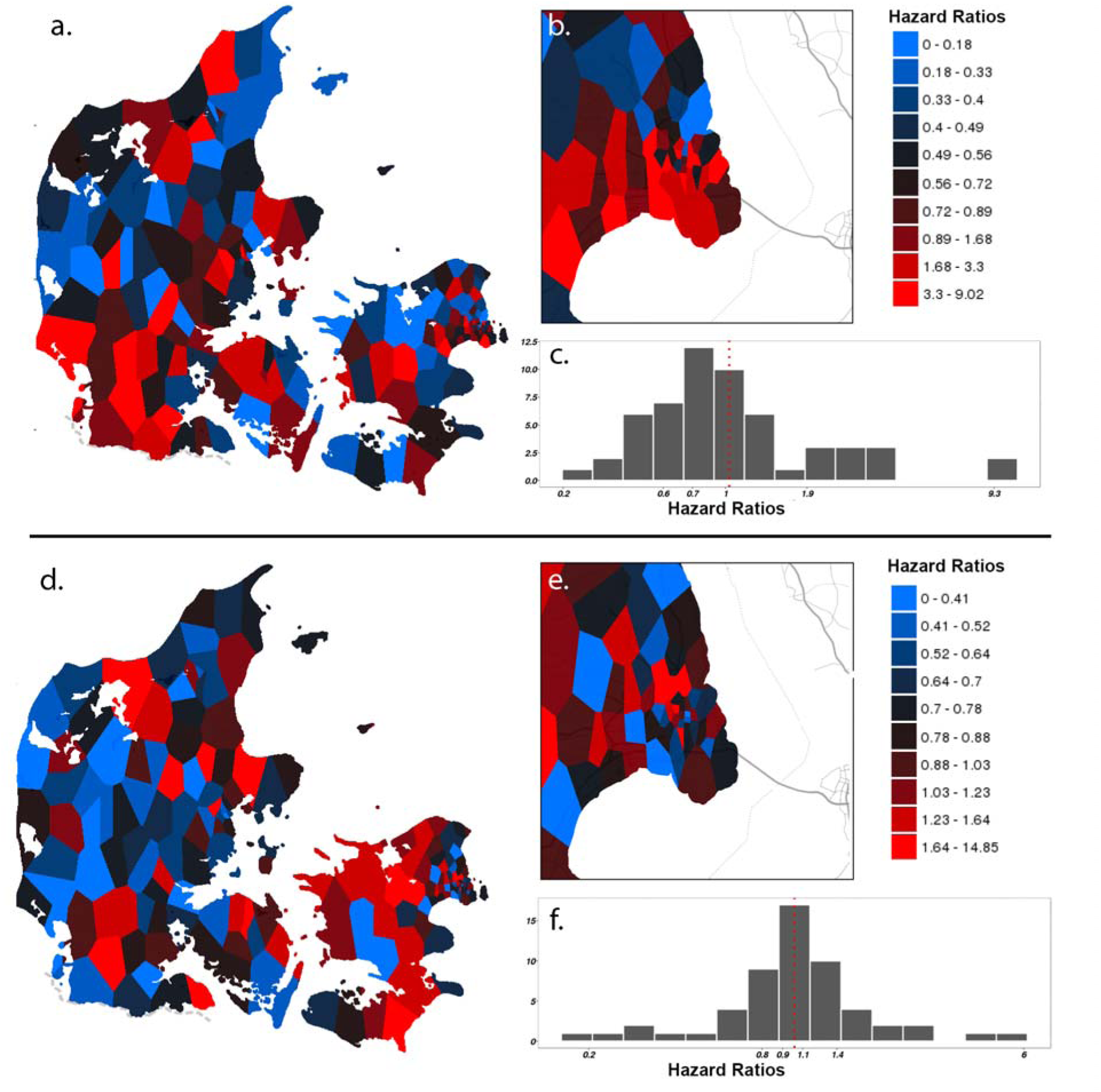
Risk distribution of E and GxE. The estimated E component is shown in the upper panel (a – c) while the estimated GxE component is hown in the lower panel (d – f). All colors were centered on national average while scaled according to risk deciles. (a) Hazard ratios distribution of E component in Denmark, (b) Hazard ratios distribution of E component in the metropolitan area, Copenhagen, (c) Histograms of E risk distribution within the metropolitan area, (d) Hazard ratios distribution of GxE component in Denmark, (e) Hazard ratios distribution of GxE component in the metropolitan area, Copenhagen, (f) Histograms of GxE risk distribution within the metropolitan area.

## Discussion

Our novel spatial mapping analysis strategy transforms the "candidate environment” approach for disease risk into a search for environmental hot spots, localizing where environmental factors appear to have a strong impact. The flexibility of this approach enables the estimation of the amount variance accounted for by E and GxE effects without direct measurement of environmental risk factors. Both simulations and empirical application demonstrate the utility of this strategy as an alternative to the candidate environment approach.

Applying this strategy to nationwide, population-based longitudinal data enriched with genetic information, we recapitulate the well-known urban-rural gradient in schizophrenia risk based on the residential information alone. Furthermore, we show that locale of upbringing significantly contributes to the risk for schizophrenia even after controlling for population density. Both E and GxE spatial effects demonstrate substantial variation within city boundaries and account for a higher proportion of schizophrenia risk than simple urban-rural contrasts. In terms of schizophrenia risk, results indicate that the locale an individual was born and raised in is more important than urban-rural differences *per se*, even within the confines of a single city. Our patterns of E and GxE across Denmark can be regarded as reference distribution. The partitioned risk contour serves as an initial guide to find the true risk element. Further comparisons with putative environmental factors can reveal the underlying elements that are highly relevant for the etiology of schizophrenia.

As a proof of concept study, our current analysis is not without limitations. First, the average age of the iPSYCH case-cohort is younger than the expected incidence peak of schizophrenia. Although the age range of our cohort is eight to thirty-two years, encompassing the incidence peak of schizophrenia, some cohort members are still at risk for schizophrenia. Right-censoring among cohort members reduces the power of statistical analyses. However, by analyzing the case-cohort with age-adjusted RR’s and survival analyses with inverse probability of sampling weights, we obtain unbiased estimates of incidence proportions. Second, our case-cohort is relatively young, while existing GWAS of schizophrenia tend to recruit more chronic patients in middle age^35^. Thus, the PRS we used may be biased toward older patients, reducing the predictive power of the already weak biological instrument. Third, as shown in our simulations, the size of the GxE effect depends upon the predictive accuracy of the G effect. Because the PRS is a weak instrument of G, the true size of the GxE effect is probably several times larger than our current estimate, as suggested by our simulations. Fourth, we did not examine the impact of migration on locale effects. Since we cannot differentiate GxE from the gene by environment correlation introduced by migration, we restricted our analyses to individuals who have Danish parents and controlled for the duration of stay at the place of birth. Although by this we intended to reduce the influence of migration, migration itself can be an important contributor for spatially-embedded risk, as many migrants tend to live in clusters, especially in urban areas. The spatial patterns we observe are unlikely due to the confounding effects of selective migration^4^ since locale of upbringing was assessed before age 7, at which age no one has yet been diagnosed with schizophrenia. Lack of selective migration is further supported by a more recent study based on the Danish registry showing no evident association between schizophrenia PRS and urbanicity ^36^.

Despite these caveats, we demonstrate that locale effects and modulating effects of locale on genetic risk account for a substantial proportion of urbanicity effects in Demark. Living in a locale with a high E component increases the risk for schizophrenia by as much as 122 percent, independent of genetic liability and family history. Meanwhile, living in a locale with a high GxE component can increase risk due to genetic liability for schizophrenia by as much as 78 percent. Because our results demonstrate risk variation with finer resolution and stronger effects than aurban-rural demarcation, there must be specific factors underlying previously observed urban effects. However, identification of factors explicating urban risk has been unsuccessful to date’^4-7^. Given the uncertainty involved, invalid constructs or measurement error could be contributors to low power to detect risk associations with specific environmental factors. Our spatial mapping strategy is an alternative approach, since finding high risk locales does not depend on correct specification of a purported environmental risk factor.

In the 19th century, epidemiology pioneer John Snow mapped high-density regions of cholera cases onto London streets and thus identified the water source as the key infectious medium. By demonstrating that the locale of upbringing significantly contributes to risk and modulates genetic susceptibility to schizophrenia, we hope this is the first step in isolating the source of spatial risk variation, facilitating the design of future public health interventions for severe mental disorders.

## Contributors

CCF, WKT, and CBP designed the study, performed data analysis, interpreted the results, and wrote the manuscript. VA, AB, and AJS collected the data, performed data quality control, and performed data analysis. JM, MJG, DG, SG, PBM, EA, and TW provide substantial inputs on revising the manuscript. All authors approved the final manuscript.

## Acknowledgments

This study was supported by the Lundbeck Foundations Initiative for Integrated Psychiatic Reseach, IPSYCH (grant numbers R102-A9118 and R155-2014-1724), Denmark, and conducted using the Danish National Biobank resource supported by the Novo Nordisk Foundation. JM was supported by a NHMRC Project John Cade Fellowship (APP1056929) and a Niels Bohr Professorship from the Danish National Research Foundation. WKT and AJS were supported by 1R01GM104400.

## Reference

1 Keller, M. C. GenexEnvironment Interaction Studies Have Not Properly Controlled for Potential Confounders: The Problem and the (Simple) Solution. Biological Psychiatry 75, 18–24, doi:10.1016/j.biopsych.2013.09.006 (2014).

2 McAllister, K. et al. Current Challenges and New Opportunities for Gene-Environment Interaction Studies of Complex Diseases. American Journal of Epidemiology 186, 753–761, doi:10.1093/aje/kwx227 (2017).

3 Pedersen, C. B. & Mortensen, P. B. Family history, place and season of birth as risk factors for schizophrenia in Denmark: a replication and reanalysis. The British Journal of Psychiatry 179, 46–-52 (2001).

4 Vassos, E., Pedersen, C. B., Murray, R. M., Collier, D. A. & Lewis, C. M. Meta-analysis of the association of urbanicity with schizophrenia. Schizophrenia bulletin 38, 1118–1123 (2012).

5 Pedersen, C. B. No evidence of time trends in the urban--rural differences in schizophrenia risk among five million people born in Denmark from 1910 to 1986. Psychological medicine 36, 211–219 (2006).

6 Pedersen, C. B. Persons with schizophrenia migrate towards urban areas due to the development of their disorder or its prodromata. Schizophrenia research 168, 204–208 (2015).

7 Pedersen, C. B. & Mortensen, P. B. Urbanization and traffic related exposures as risk factors for schizophrenia. BMC psychiatry 6, 2 (2006).

8 Brown, A. S. & Derkits, E. J. Prenatal infection and schizophrenia: a review of epidemiologic and translational studies. American Journal of Psychiatry 167, 261–280 (2009).

9 Werner, S., Malaspina, D. & Rabinowitz, J. Socioeconomic Status at Birth Is Associated With Risk of Schizophrenia: Population-Based Multilevel Study. Schizophrenia Bulletin 33, 1373 (2007).

10 van Os, J., Hanssen, M., Bak, M., Bijl, R. V. & Vollebergh, W. Do Urbanicity and Familial Liability Coparticipate in Causing Psychosis? American Journal of Psychiatry 160, 477–482 (2003).

11 van Os, J., Pedersen, C. B. & Mortensen, P. B. Confirmation of Synergy Between Urbanicity and Familial Liability in the Causation of Psychosis. American Journal of Psychiatry 161, 2312–2314 (2004).

12 Krabbendam, L. & Van Os, J. Schizophrenia and urbanicity: a major environmental influence---conditional on genetic risk. Schizophrenia bulletin 31, 795–799 (2005).

13 Grech, A., van Os, J. & Investigators, G. Evidence That the Urban Environment Moderates the Level of Familial Clustering of Positive Psychotic Symptoms. Schizophrenia Bulletin 43, 325–331, doi:10.1093/schbul/sbw186 (2017).

14 Visscher, P. M. et al. 10 Years of GWAS Discovery: Biology, Function, and Translation. The American Journal of Human Genetics 101, 5–22, doi:https://doi.org/10.1016/j.ajhg.2017.06.005 (2017).

15 Snow, J. On the mode of communication of cholera. (John Churchill, 1855).

16 Kelsall, J. & Wakefield, J. Modeling spatial variation in disease risk: a geostatistical approach. Journal of the American Statistical Association 97, 692–701 (2002).

17 Faris, R. E. L., & Dunham, H. W. Mental disorders in urban areas: an ecological study of schizophrenia and other psychoses. (Univ. Chicago Press, 1939).

18 Barr, C. D. & Schoenberg, F. P. On the Voronoi estimator for the intensity of an inhomogeneous planar Poisson process. Biometrika, 977–984 (2010).

19 Pedersen, C. B. et al. The iPSYCH2012 case-cohort sample: new directions for unravelling genetic and environmental architectures of severe mental disorders. Mol Psychiatry (2017).

20 Price, A. L. et al. Principal components analysis corrects for stratification in genome-wide association studies. Nature genetics 38, 904 (2006).

21 Abraham, G., Qiu, Y. & Inouye, M. FlashPCA2: principal component analysis of biobank-scale genotype datasets. bioRxiv (2016).

22 O’Connell, J. et al. Haplotype estimation for biobank-scale data sets. (2016).

23 Howie, B. N., Donnelly, P. & Marchini, J. A flexible and accurate genotype imputation method for the next generation of genome-wide association studies. PLoS Genet 5, e1000529 (2009).

24 Consortium, G. P. & others. An integrated map of genetic variation from 1,092 human genomes. Nature 491, 56 (2012).

25 Ripke, S. et al. Biological insights from 108 schizophrenia-associated genetic loci. Nature 511, 421 (2014).

26 Team, R. C. R: A Language and Environment for Statistical Computing. (2016).

27 Baddeley, A. & Turner, R. spatstat: An R Package for Analyzing Spatial Point Patterns. Journal of Statistical Software 12, 1–42 (2005).

28 Therneau, T. M. coxme: Mixed Effects Cox Models. (2018).

29 Kahle, D. & Wickham, H. ggmap: Spatial Visualization with ggplot2. The R Journal 5, 144–161 (2013).

30 Cheng, J., Karambelkar, B. & Xie, Y. leaflet: Create Interactive Web Maps with the JavaScript ‘Leaflet’ Library. (2017).

31 Chang, W., Cheng, J., Allaire, J. J., Xie, Y. & McPherson, J. shiny: Web Application Framework for R. (2017).

32 Rothman, K. J., Greenland, S. & Lash, T. L. Modern epidemiology. (2008).

33 Barlow, W. E., Ichikawa, L., Rosner, D. & Izumi, S. Analysis of Case-Cohort Designs. Journal of Clinical Epidemiology 52, 1165–1172, doi:10.1016/S0895-4356(99)00102-X (1999).

34 Austin, P. C., Wagner, P. & Merlo, J. The median hazard ratio: a useful measure of variance and general contextual effects in multilevel survival analysis. Stat Med 36, 928–938, doi:10.1002/sim.7188 (2017).

35 Meier, S. M. et al. High loading of polygenic risk in cases with chronic schizophrenia. Mol Psychiatry 21, 969–974 (2016).

36 Paksarian, D. et al. The role of genetic liability in the association of urbanicity at birth and during upbringing with schizophrenia in Denmark. Psychological Medicine, 1–10 (2017).

